# Plasma triacylglycerol length and saturation level mark healthy aging groups in humans

**DOI:** 10.1101/2024.08.22.609162

**Authors:** Weisha Li, Bauke V. Schomakers, Michel van Weeghel, Lotte Grevendonk, Frédéric M. Vaz, Gajja S. Salomons, Patrick Schrauwen, Joris Hoeks, Arwen W. Gao, Riekelt H. Houtkooper, Georges E. Janssens

## Abstract

Complex lipids, essential components in biological processes, exhibit conserved age-related changes that alter membrane properties, cellular functions, and are implicated as biomarkers and contributors to longevity and age-related diseases. While physical activity alleviates age-related comorbidities and physical impairments, comprehensive exploration of the underlying biological mechanisms, particularly at the level of complex lipids, remains limited. However, clinical studies suggest that physical activity may counteract these age-related lipidomic changes, presenting a promising avenue for intervention. We performed lipidomic profiling of plasma from an extensively characterized cohort of young and aged individuals. Annotating 1446 unique lipid species across 24 lipid classes we found the most prominent difference in older adults was an accumulation of triacylglycerols (TGs), with lower physical activity levels associated with higher TG levels in plasma and reduced physical functionality. Remarkably, lipid species in the TG class did not accumulate uniformly. Rather, our study unveiled a negative correlation between higher physical activity levels and TGs with shorter chain length and more double bonds in this demographic. Overall, our research highlights that plasma TG length and saturation level can help mark healthy aging groups in humans. These findings deepen our understanding of how aging affects complex lipids, and the influence of physical activity on this process.

## Introduction

Aging is an inherent biological phenomenon characterized by progressive changes in various physiological processes that ultimately result in a decline in general health and functionality. Multiple key hallmarks have been recognized and delineated as contributing to the aging process. These twelve hallmarks include, amongst others, mitochondrial dysfunction, impaired immune function, reduced autophagy, epigenetic alterations, aberrant intercellular communication, loss of telomeres, altered nutrient sensing, and impaired protein homeostasis [1–6]. Understanding the drivers of different rates of biological aging could enable the development of more tailored and effective preventive and therapeutic strategies aimed at maximizing health span [7]. This understanding could also enhance diagnostics of aging, including more accurate biological clocks and providing a clearer assessment of aging [8, 9]. The determinants of biological aging remain largely unknown, yet lipid metabolism has been suggested to be involved [10–12].

Lipids are very diverse and complex biological compounds that play crucial roles in a variety of biological processes [13]. Indeed, age-related changes in complex lipids are linked to longevity and age-related diseases, serving as both biomarkers and contributors [12, 14–16]. Therefore, plasma lipid profiles hold promise as biomarkers for age-related changes and associated disease risks [15, 17, 18]. Certain lipid species in human plasma, such as ceramides (CERs), sphingomyelins (SMs), lysophosphatidylcholines (LPCs), and cholesteryl esters (CEs), tend to increase with age [15, 19]. Studies have also indicated higher levels of certain sphingomyelin (SM) species such as SM(d18:1/18:2) and SM(d18:1/17:0), positively impact healthy aging [20]. Moreover, the concentration of plasma long-chain free fatty acids is inversely correlated with longevity across various mammalian species, suggesting their potential role as a biomarker for longevity [18]. Changes in plasma triacylglycerol (TG) levels and composition also occur with age [21], four long-chain and highly polyunsaturated TG species, TG(54:7), TG(54:6), TG(56:7), TG(56:6), are positively correlated to human longevity in the plasma lipidome [22, 23]. Additionally, factors related to lipid changes have been shown to regulate longevity [12, 18, 23]. For example, different types of intermittent fasting and caloric restriction can significantly improve the serum lipid profile by reducing TGs, total cholesterol and low-density lipoprotein cholesterol concentrations, which are associated with age-related diseases [24–26].

Furthermore, regular physical activity is associated with improved plasma lipid profiles in the elderly, including lower levels of TGs, total cholesterol, and LDL cholesterol, as well as higher levels of HDL cholesterol [27–30]. Physical activity is therefore also associated with reduce cardiovascular risks and mortality in older people [31–33]. However, the understanding of how lipidomic patterns in the elderly are influenced by different physical activity status remains unclear. Understanding inter-individual differences in lipid characteristics and the effect of physical activity levels is a promising avenue of research.

We performed lipidomic profiling of plasma from an extensively characterized cohort of young and aged individuals, where the older adults consisted of both very physically trained and untrained individuals [34, 35]. Our investigation revealed TG as the predominant lipid class accumulated in the plasma of older adults. Notably, our findings indicated a health-related trend associated with TG levels in older adults, indicating a correlation between a reduction in TG length and double bonds, coinciding with a decline in physical functionality. Our research highlights that plasma TG length and saturation level can help differentiate healthy aging groups in humans. These findings deepen our understanding of aging processes at the lipidomic level, emphasizing the intricate relationship between physical activity levels and lipid metabolism in aging.

## Results

### The lipidome of aging human plasma

To better understand which lipids are associated with aging and how physical active levels relate to plasma lipid aging we performed semi-targeted lipidomic profiling of plasma from an extensively characterized cohort of young and older individuals (Figure 1A, Table S1), which we previously established (clinicaltrails.gov identifier NCT03666013)[34, 35]. The specific cohort analyzed here included 14 young individuals (young adults) whereas the group of older participants consisted of exercise-trained older adults (trained older adults, n=19), older adults with normal physical activity levels (normal older adults, n=16), and physically impaired older adults (impaired older adults, n=3)) [36]. All participants were free of any diseases. Young and normal older adults took an average of ∼10000 steps per day and spent <1 hour of their active time in high-intensity activities (Table S1). The trained older adults were more active (∼13000 steps per day on average) and had more time spent in high-intensity activities (>3h), whereas the physically impaired were less active (∼6000 steps per day on average) and had less time spent in high-intensity activities (<1h).

**Figure 1.**
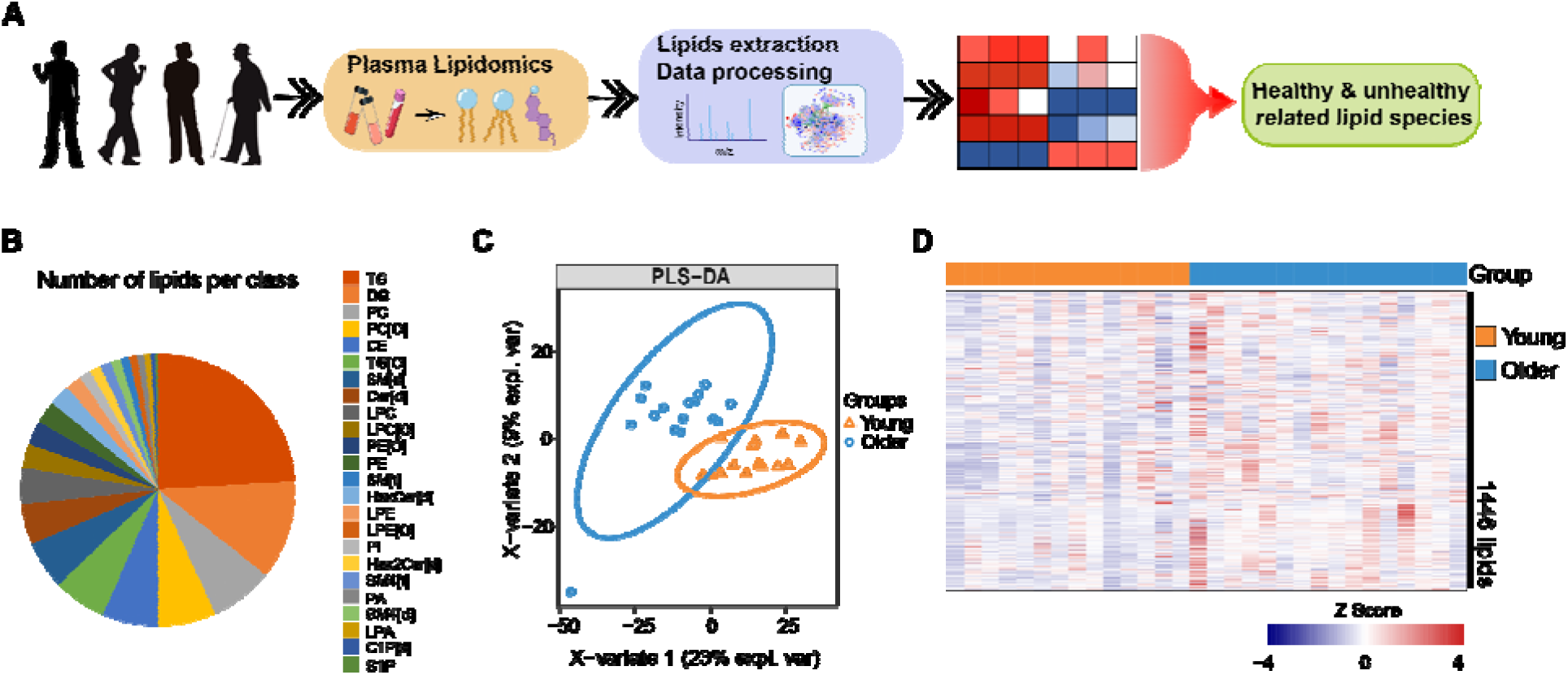
The lipidome of aging human plasma. (**A**) Flowchart depicting the inclusion of study participants categorized as either young or older adults, with the subsequent acquisition of plasma samples for mass spectrometry-based lipidomic analysis. (**B**) Pie chart showing the number of lipids of each lipid class as a percentage of the total number of detected lipids in plasma. (**C**) Partial least squares discriminant analysis (PLS-DA) of lipidomes of young and older individuals possessing different lipids levels (young versus normal older adults. young individuals: n=14; normal older adults: n=16). (**D**) The heatmap illustrates a comparative analysis of lipid profiles between young adults and normal older adults, showcasing variations in lipid abundance. Elevated levels are represented in red, while reduced levels are indicated in blue.

Using ultrahigh-performance liquid chromatography coupled to a high-resolution mass spectrometer, we conducted a comprehensive analysis of plasma samples obtained from both young and older adults of the three different physical fitness levels (Figure 1A, Table S2). We annotated 1446 unique lipid species distributed across 24 lipid classes (Figure 1B, Table S2). To better understand the lipid profile changes that occur with aging, we performed a Partial Least Squares Discriminant Analysis (PLS-DA) on all groups, which showed a general separation between young individuals and normal older adults (Figure 1C), suggesting aging produces significant changes on plasma lipids. To explore the biological differences between young and older groups, we compared the lipid abundance between young individuals and older adults. Overall, we found a general increase in the levels of multiple lipids in older adults (Figure 1D). Together, these plasma lipidomic findings illustrate the distinction between young individuals and older adults, highlighting an age-associated increase in most lipids.

### Prevalent accumulation of triacylglycerol (TG) species among older adults versus young individuals

We next analyzed the lipid contribution similarities and differences in the plasma of the three groups of older adults with varying physical activity levels by comparing the lipids that are more or less abundant. Here we found age-associated increase of lipids in all three groups of older adults (Figure 2A-C). Since only a few lipids significantly depleted in the age groups compared to young adults, we shifted our focus to those that increased. Within this set of significantly increased lipids, 54 lipids were significantly higher in older adults regardless of their activity status (Figure 2D). Of these, 48 were identified as TG species, five diacylglycerol (DG) species, and one phosphatidylinositol (PI) species (Figure 2D-E).

**Figure 2.**
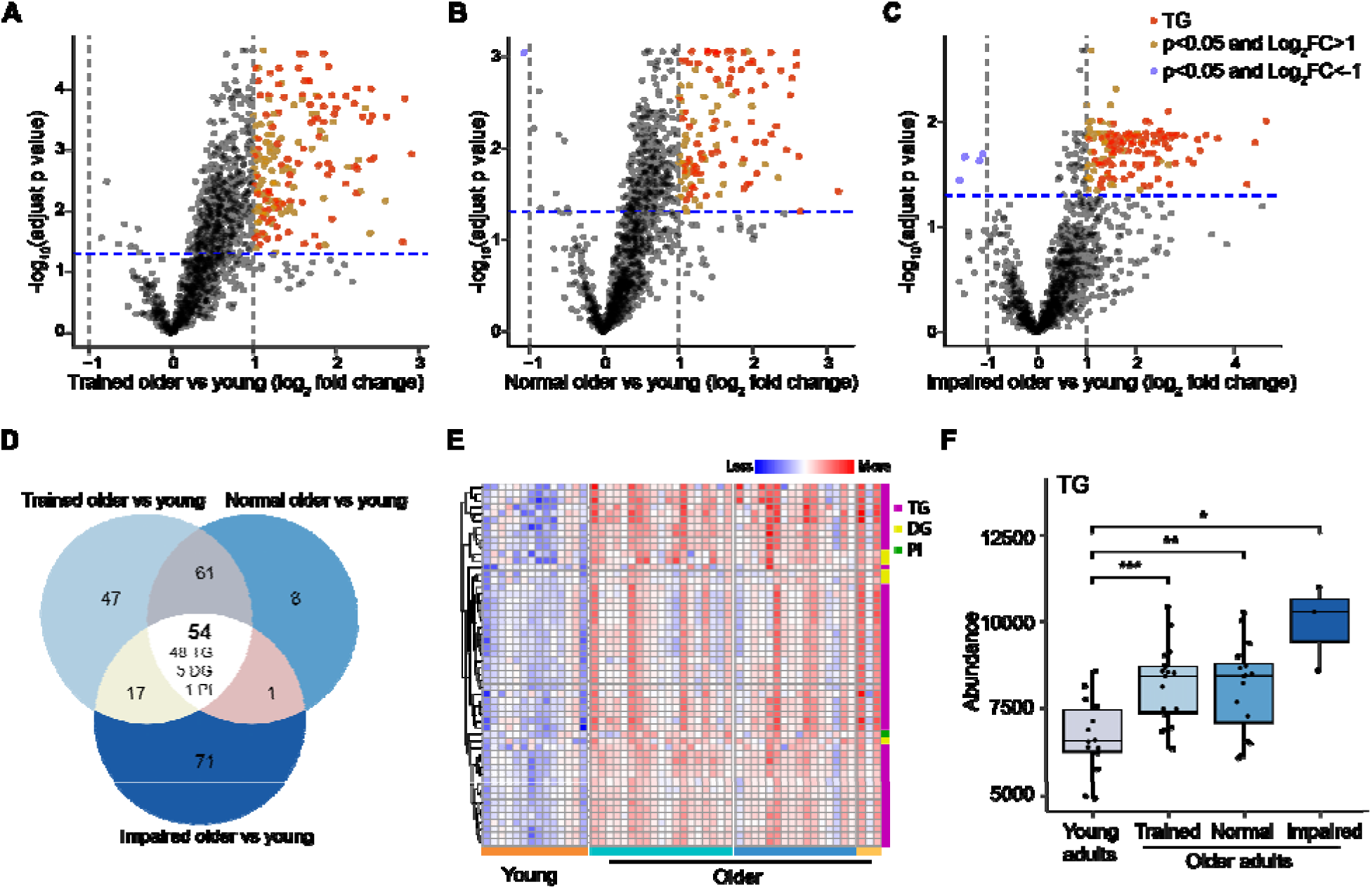
Prevalent accumulation of TG species among elderly versus young individuals. (**A-C**) Volcano plot of fold change (x-axis, log_2_ scale) versus adjusted p-value (y-axis, -log_10_ scale) for impaired older adults. Values on x-axis greater than zero indicate accumulation with age, less than zero indicates depletion with age. (A), normal older adults (B), or trained older adults compared (C) to young individuals, illustrating significant depleted(blue) or accumulated (red) lipids. The horizontal line indicates significance (adjusted p-value <0.05), and the vertical line shows absolute log_2_ fold change >1. Significance was determined using an empirical Bayes-moderated t-test. (**D**) Venn diagram of the overlap of significantly higher (red in panels A-C) abundance lipids in each aged group (impaired older adults, normal older adults and trained older adults) compared to young individuals (p < 0.05 and absolute log_2_ fold change >1). (**E**) Heatmap of 54 lipids that accumulated in impaired, normal and trained older adults compared to young adults. (**F**) Abundance of TGs in the four groups (young individuals, trained older adults, normal older adults, and impaired older adults). Sample sizes: young individuals: n = 14; trained older adults: n = 19; normal older adults: n = 16; impaired older adults: n = 3. Significance was determined using an unpaired Student’s t-test, *: p<0.05, **: p<0.01, ***: p<0.001.

When investigating the total levels of lipid classes, we found that TG was not only one of the most accumulated lipid classes in healthy older adults when compared to young individuals but also displayed an association with activity levels, with more severe TG accumulation in impaired older adults (Figure 2F). These findings are in line with numerous studies that have consistently reported plasma TG levels to be higher in older adults versus younger adults [37–40]. Therefore, with our lipidomics offering a lens to not only focus on TG abundance but also on chain length and saturation levels, we found it of interest to further explore how TG subclasses significantly accumulate in the young compared to older adults with varying physical activity levels.

### TG carbon chain length and saturation levels have different distributions in older adults compared to young individuals

To further investigate how subsets of TG species change with age, we investigated their composition further. Carbon chain length and the number of double bonds in those chains displayed divergent patterns (Figure 3A-C). For instance, TG in trained older adults showed markedly higher levels of highly unsaturated lipids when compared to young individuals (>4 double bonds) (Figure 3A). Similarly, changes in the TG composition of normal older versus young also showed a significant increase of TG with highly unsaturated double bonds (>4 double bonds) (Figure 3B). Conversely, impaired older adults exhibited a significant increase in lipids representing a broader range of double bonds, with increases in polyunsaturated fatty acid (PUFA)-containing species and also in species with fewer double bonds (≤4 double bonds) (Figure 3C). Indeed, impaired older adults exhibited a high level of TGs with shorter carbon-chain lengths (<56) in comparison to both normal older adults and trained older adults (Figure 3D-E). The composition of aged groups compared to each other showed that both normal and trained older adults differed from impaired older adults in a highly similar way (Figure 3D-E). Together, our data suggest that TGs with fewer double bonds (≤ 4 double bonds) and shorter carbon-chain lengths (<56) accumulated more in impaired older adults, whereas trained older adults have more extremely unsaturated TG species.

**Figure 3.**
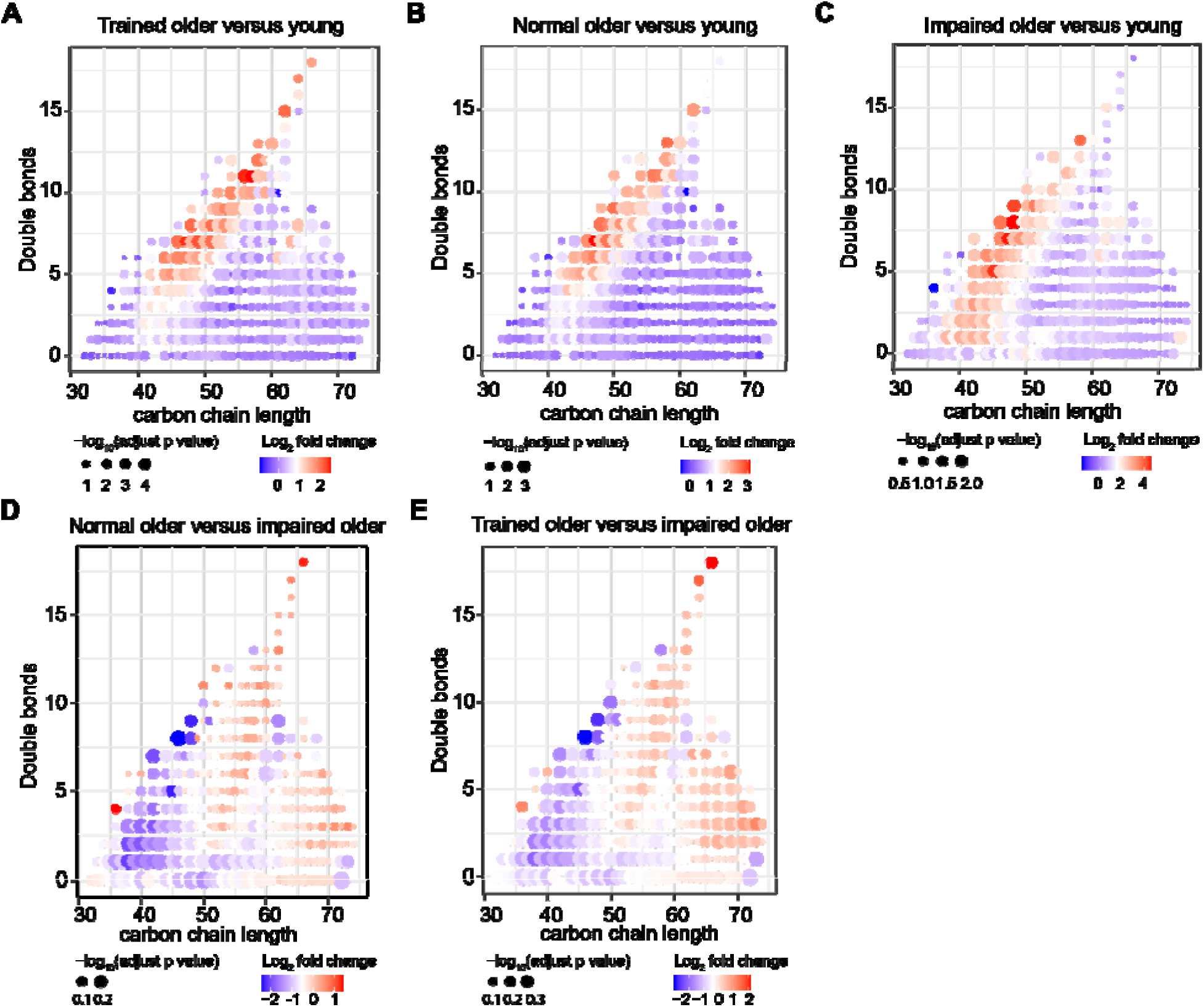
TG carbon chain length and saturation levels have different distributions in older adults compared to young adults. (**A**) Changes in the TG composition of trained older versus young show a significant increase of phospholipids with shorter carbon-chain length (<56) and phospholipids with highly polyunsaturated double bonds (>4 double bonds). The log_2_ fold change and adjusted p-value were calculated by comparing trained older adults with young individuals. The red color indicates lipids accumulated more in trained older adults. (**B**) Changes in the TG composition of normal older versus young show a significant increase of phospholipids with highly polyunsaturated double bonds (>4 double bonds) and a significant decrease of lipids with ≤4 double bonds and phospholipids with longer carbon-chain length (≥56). The log_2_ fold change and adjusted p-value were calculated by comparing normal older adults with young individuals. The red color indicates lipids accumulated more in normal older adults. (**C**) Changes in the TG composition of impaired older versus young show a significant decrease of phospholipids with longer carbon-chain length (≥56) and lipids with highly polyunsaturated double bonds (>4 double bonds). The log_2_ fold change and adjusted p-value were calculated by comparing impaired older adults with young individuals. The red color indicates lipids accumulated more in impaired older adults. (**D-E**) Changes in the TG composition of normal older adults (D) and trained older adults (**E**) versus impaired older adults show a significant increase of phospholipids with longer carbon-chain length in normal and trained older (≥56). The log_2_ fold change and adjusted p-value were calculated by comparing normal (D) and trained (E) older adults to impaired older adults. The red color indicates lipids accumulated more in normal older adults(D) and trained older adults(E). Sample sizes are n = 14 for young individuals, n = 19 for trained older adults, n = 16 for normal older adults and n = 3 for impaired older adults.

### Physical activity level is negatively correlated with carbon chain length and saturation levels in older adults

To further explore the relations between TG composition and physical activity statuses, we classified the TGs into 6 subgroups based on the saturation level and carbon chain length as follows: (a) “shorter-saturated” had the criteria of TG carbon chain length < 56 and double bonds = 0; (b) “shorter-less-unsaturated” had the criteria of carbon chain length < 56 and double bonds ≤ 4; (c) “shorter-more-unsaturated” had the criteria of carbon chain length < 56 and double bonds >4; (d) “longer-saturated” had the criteria of carbon chain length ≥ 56 and double bonds = 0; (e) “longer-less-unsaturated” had the criteria of carbon chain length ≥ 56 and double bonds ≤ 4; (f) “longer-more-unsaturated” had the criteria of carbon chain length ≥ 56 and double bonds >4 (Figure 4A).

**Figure 4.**
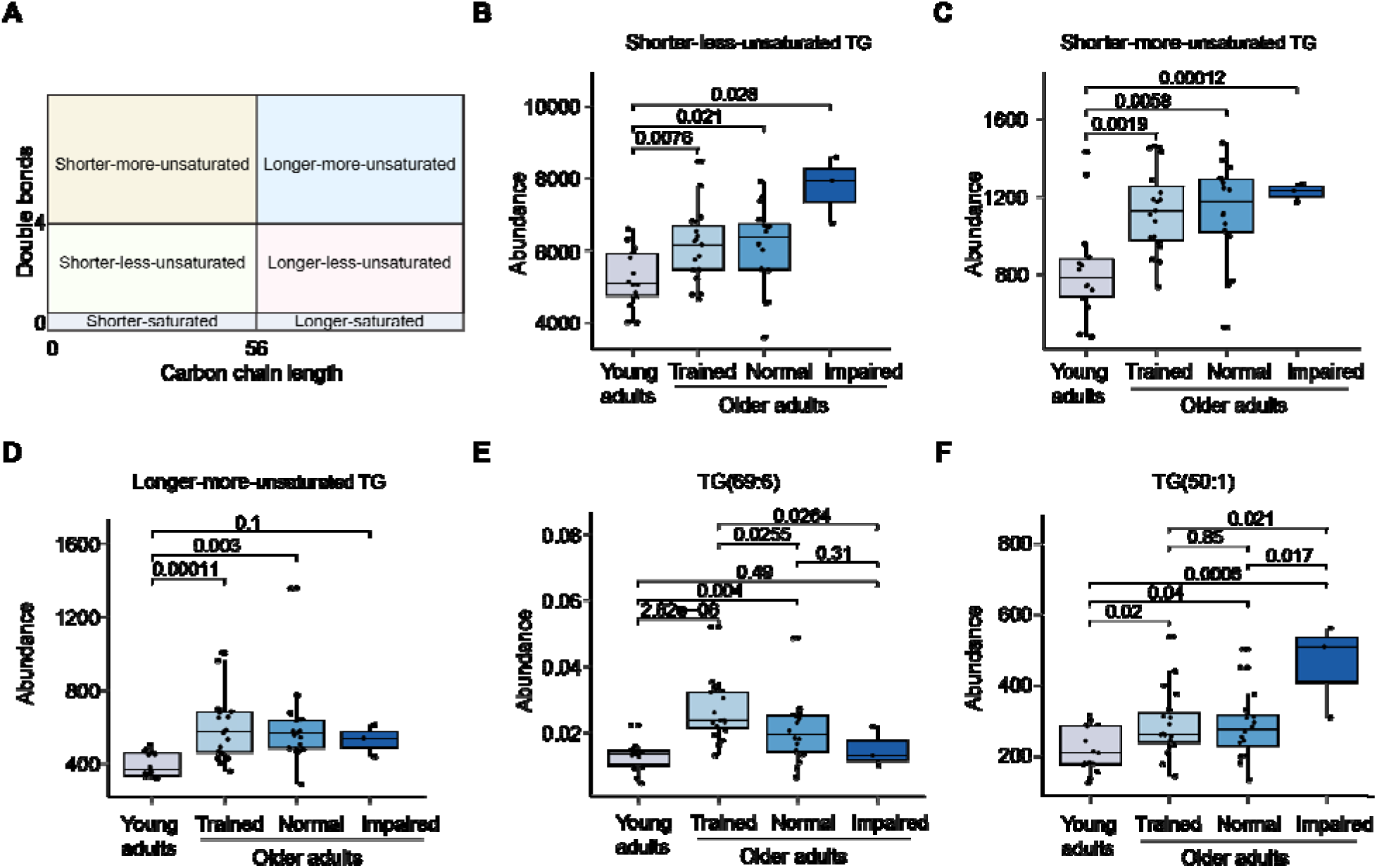
Physical activity level is negatively correlated with carbon chain length and saturation levels in older adults. (**A**) Classification of TGs into six categories based on carbon chain length and double bonds level, segregated at thresholds of 56 to distinguish between longer chain length and shorter chain length, and at 0 and 4 to delineate saturated and more or less unsaturated. (**B-D**) Abundance of shorter-less-unsaturated TG (B), shorter-more-unsaturated TG (C), and longer-more-unsaturated TG (D) in the four groups (young individuals, trained older adults, normal older adults, and impaired older adults). Significance was determined using an unpaired Student’s t-test. (**E-F**) Abundance of TG(50:1) (E) and TG(69:6) (F) in the four groups (young individuals, trained older adults, normal older adults, and impaired older adults). Significance was determined using an empirical Bayes-moderated t-test. Sample sizes are n = 14 for young individuals, n = 19 for trained older adults, n = 16 for normal older adults, and n = 3 for impaired older adults.

Then, we conducted a comparative analysis of TG subgroup abundance levels within the different groups (Figure 4B-D). Our findings revealed that impaired older adults exhibited higher levels of shorter-less-unsaturated TGs (Figure 4B), reflecting the previous pattern we identified of total TGs (Figure 2F). Furthermore, the three groups of older adults all displayed a higher level of shorter-more-unsaturated TGs (Figure 4C). Meanwhile, longer-more-unsaturated TGs were more accumulated in the trained older adults (Figure 4D). These data demonstrate that the level of physical activity in older adults relates to levels of TG chain length and double bonds, where impaired older adults have a greater abundance of shorter-less-unsaturated TGs and trained older adults possess a greater abundance of longer-more-unsaturated TGs. Finally, specific lipid species display similar trends in differentiating between the older adults with varying physical activity levels. For example, the low abundant TG(69:6), a TG containing very long-chain fatty acids, was most abundant in the trained older adults (Figure 4E), while in contrast TG(50:1), a shorter TG, was highest in the impaired older adults (Figure 4F).

### Association between physiological phenotypes and TG composition in older adults

To further delineate how physiological phenotypes of our study participants were related to the TG distributions, we performed a cross-correlation of the six categories of TGs (Figure 4A) to physical parameters in older individuals (Figure 5A). Surprisingly, body mass index (BMI) showed a positive correlation with total TG, particularly shorter-less-unsaturated TGs, which accumulate more in physically impaired older adults. However, there was no significant correlation between BMI and more-unsaturated TGs, nor was there a significant correlation between BMI and physical activity levels (movement level of adults) (Figure 5A). However, we found that movement level (steps per day) showed a distinct negative correlation with the shorter-more-unsaturated TGs (R = -0.38, Spearman’s *p = 0.022*) (Figure 5A-B), while correlations with other TG subsets were not significant. Similarly, whole-body insulin sensitivity (measured as glucose infusion rate during a glucose clamp) had a strong negative correlation with the shorter-more-unsaturated TGs (R = 0.39, Spearman’s *p = 0.017*) (Figure 5A, C). Furthermore, we found a positive correlation between diastolic and systolic blood pressure and shorter-more-unsaturated TGs (R = 0.39, Spearman’s *p = 0.014* and R = 0.32, Spearman’s *p = 0.05* respectively) (Figure 5A and 5D-E).

**Figure 5.**
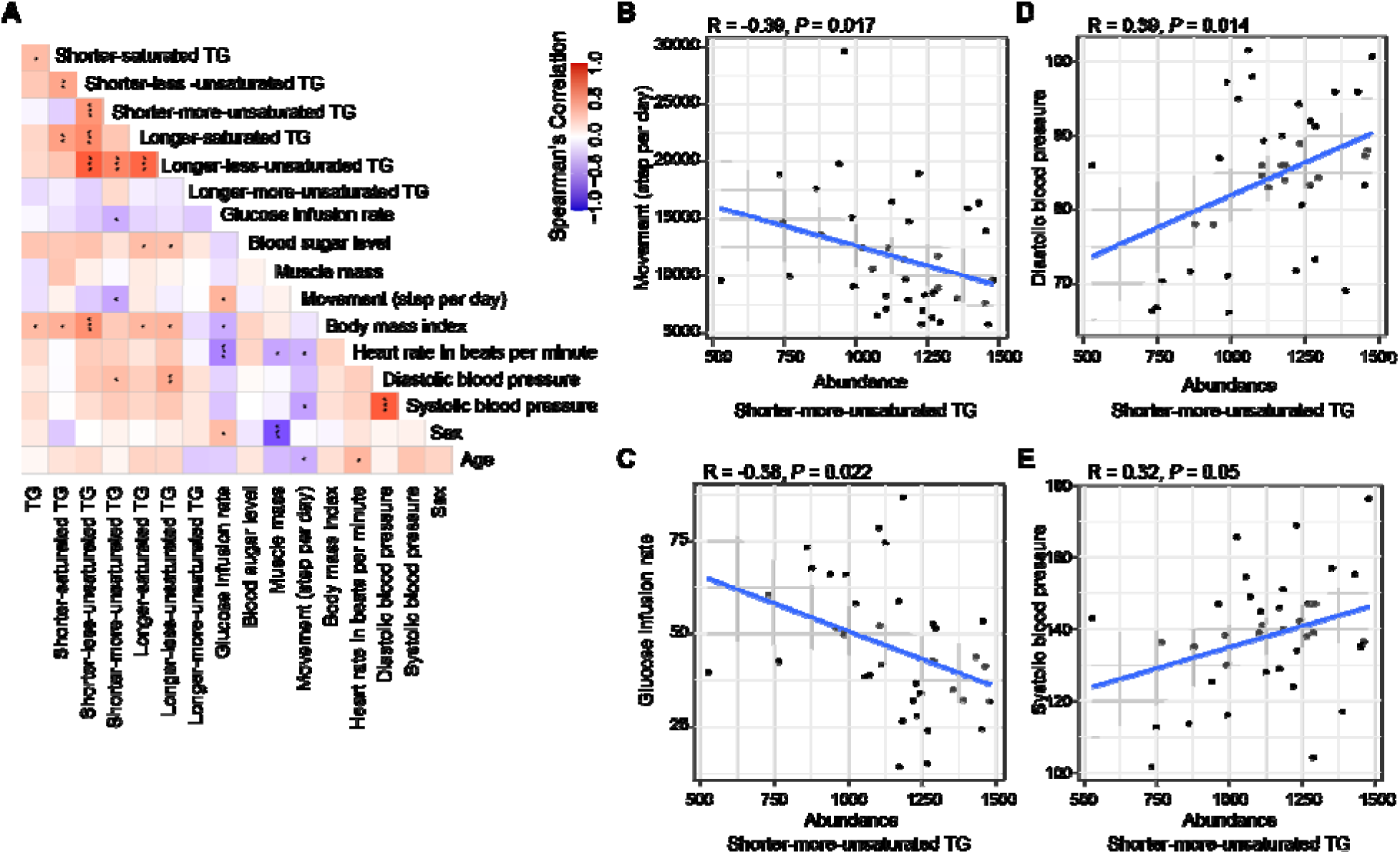
Association between physiological phenotypes and TG composition in older adults. (**A**) Spearman’s product-moment correlation coefficient matrix illustrates associations between TG levels and physical conditions in a cohort of four old groups (n = 38). Color scale: blue (negative correlation) to white (no correlation) to red (positive correlation). (**B-E**) Scatterplots illustrating specific correlations between (B) average daily steps and shorter-more-unsaturated TG, (C) glucose infusion rate and shorter-more-unsaturated TG, (D) diastolic blood pressure and shorter-more-unsaturated TG, (C) systolic blood pressure and shorter-more-unsaturated TG among the elderly (n = 38). Statistical significance was assessed using Spearman’s rank correlation coefficient. *: p<0.05, **: p<0.01, ***: p<0.001.

Taken together, we here present a view of the strong relationship between the shorter-more-unsaturated TGs and movement levels in older adults, and our study also revealed a relationship between shorter-more-unsaturated TGs and systolic and diastolic blood pressure and glucose infusion rate.

## Discussion

In the current study, we investigated age-related plasma lipidomic changes by comparing young and older individuals. To identify how plasma TG differences in older adults relate to physical activity, we evaluated three groups of older adults ranging in physical fitness. Our investigation revealed that (1) TGs accumulated with age regardless of more or less physical activity, though (2) older adults with lower physical activity levels show much higher TG levels and finally (3) plasma triglyceride length and saturation level can help distinguish healthy aging groups in humans.

Although age is clearly a risk factor for metabolic disease, the mechanisms that contribute to this age-related risk have not been fully elucidated. One major contributing factor to metabolic disease is aberrant TG metabolism [41, 42]. Individuals with metabolic syndrome or type 2 diabetes (T2D) often exhibit elevated TG content in the heart and liver [42]. Furthermore, altered plasma TG metabolism plays a crucial role in the development of metabolic diseases [41]. Additionally, high levels of plasma TG serve as a risk factor for cardiovascular disease [43]. Many studies have confirmed that plasma TG levels are higher in older adults versus younger adults [21, 44, 45]. However, accumulation of TGs can also occur in athletes [46, 47], therefore understanding how TG lipid composition relates to health is of paramount importance. Considering that double bonds and carbon chain length are crucial features distinguishing lipids from general metabolites, our research delves deeply into the relationship between TG length, saturation level, and aging. A more comprehensive understanding of plasma TG’s role should facilitate the development of markers for risk prediction, diagnosis, prognosis, and response to therapies, potentially guiding new directions for therapeutic interventions. In line with this, our study demonstrated an association between shorter-more-unsaturated TGs, and systolic and diastolic blood pressure, as well as insulin sensitivity.

According to the TG chain length and double bond distribution, we defined six TG subgroups and found a negative relationship between the shorter-more-unsaturated TGs and physical fitness level in older individuals. Higher blood TG is an indication of cardiovascular disease, which is among the leading causes of death worldwide [48, 49].

Considerable evidence supports physical activity as an effective adjunctive intervention for abnormal lipid levels, with studies revealing that regular exercise can help lower TG levels in the blood [50–52]. Our study demonstrates that physical activity correlates with plasma TGs in older adults, particularly those with specific chain lengths and saturation levels, which are commonly elevated in older adults. Previous research has reported that longer lifespans in various species were correlated with a higher average fatty acid chain length in the plasma [53]. Conversely, shorter-less-unsaturated TG content was reported to be associated with an increased risk of metabolic disorders like type 2 diabetes, implying their accumulation during decreased metabolic activity [54]. This aligns with our study, which found that impaired older adults accumulate more shorter-less-unsaturated TGs. This suggests that carbon chain length and the number of double bonds also play a major role in healthy aging.

Studies have observed a moderate positive association between physical activity and circulating levels of behenic acid (22:0) and lignoceric acid (24:0), which are considered to be very long-chain fatty acids (VLCFAs) [55, 56]. This association suggests that physical activity might enhance the synthesis or availability of these fatty acids, although the precise mechanism remains unclear [55]. Physical activity has been shown to affect the expression and activity of enzymes involved in fatty acid metabolism [57, 58]. For instance, physical activity can reduce the activity of stearoyl-CoA desaturase (SCD1) in adipose tissue, which is involved in the synthesis of monounsaturated fatty acids [58]. This modulation could potentially explain the increased longer-more-unsaturated TGs in the trained older adults.

Several limitations exist in our study. For example, variation may exist within the defined health groups, and the group-based analyses may have a lower resolution to detect differences than an ungrouped approach. Additionally, a limitation arises from the small sample size of impaired older adults (n=3), potentially impacting statistical analyses across distinct health groups. This small sample size also makes it virtually impossible to stratify the results across genders. Finally, our investigation into the correlation between physical activity and TG levels is not founded on data collected through intentional exercise interventions imposed on subjects. Rather, it relies on analyzing participants’ health statuses and daily physical activities. Consequently, to delineate the precise influence of aerobic and anaerobic exercise on TG levels and composition with the health trend, a clinical trial with exercise would be warranted.

Some of our reasons for conducting plasma lipidomics were to identify biomarkers, candidate causal influencers in aging, and to provide direction for future research. Our study provided evidence that aging and physical activity levels relate to TG carbon chain length and the number of carbon bonds. shorter-more-unsaturated TGs are more prevalent in older adults, potentially serving as markers of aging. Specifically, TG species such as TG(54:7), TG(54:6), TG(56:7), and TG(56:6), previously identified as markers of aging [23], are also included in this group. Moreover, shorter-more-unsaturated TGs appear to be associated with healthier aging, whereas shorter-less-unsaturated TGs are more prevalent among less-healthy older adults. This suggests the clinical utility of assessing TG characteristics in marking health status among older individuals. Exploring the effects of different types of physical exercises on lipid profiles within these demographic groups will yield valuable insights. Furthermore, expanding the sample size to include individuals with diverse health statuses would enhance comprehension of the changes in TG carbon chain length and carbon bond numbers in response to health trends.

Recent studies have demonstrated that in regression models using “-omic” platforms to predict chronological age, the residual variation in predicted age correlates with health outcomes [59–61]. These findings suggest that “omic clocks” can serve as measures of biological age. Key metabolites, such as kynurenine, phenylalanine, and those involved in steroid hormone biosynthesis, have been identified for their high predictive accuracy in predicting age [60]. Additionally, consistent age-related changes in metabolites like pyruvate, glutathione, ATP, and NAD^+^ highlight their potential as global aging markers [36, 62]. Integrating metabolic aging clocks with DNA methylation and transcriptomic clocks could provide a more comprehensive measure of biological age [63–65]. Similarly, bis(monoacylglycerol)phosphate (BMP) and sphingomyelins, phosphocholines, ceramides, and fatty acids have been recognized as useful components of lipidomic aging clocks [12, 66, 67]. Lipidomic markers are advantageous as they reflect functional aging processes, are relatively accessible to measure, and are dynamically modulated by interventions targeting aging [68]. However, challenges including standardizing lipidomics platforms, accounting for confounders, validating findings across diverse populations, and identifying causal roles of specific lipids still remain. Overall, our findings implicate TG species subsets in healthy aging biology and future work integrating these lipid-based markers into comprehensive aging clocks may serve to enhance precision measurements of healthy aging in the clinic.

## Methods

### Study participants

Fifty-two participants, including 14 young (7 male and 7 female) and 38 older (21 male and 17 female) individuals, were recruited in the community of Maastricht and its surroundings through advertisements at Maastricht University, local newspapers, supermarkets, and sports clubs. The study protocol was approved by the institutional Medical Ethical Committee and conducted in accordance with the declaration of Helsinki. All participants provided written informed consent and the study was registered at clinicaltrials.gov with identifier NCT03666013.

### Human subjects and procedures

Participants were categorized into the following study groups: young individuals with normal physical activity (20–30 years), older adults with normal physical activity (65–80 years), trained older adults (65–80 years) and physically impaired older adults (65–80 years). Participants were considered normal physically active if they completed no more than one structured exercise session per week. Participants were considered trained if they engaged in at least three structured exercise sessions of at least 1 h each per week for an uninterrupted period of at least 1 year. Participants were classified as older adults with impaired physical function in case of an SPPB score of ≤9. The SPPB score was calculated according to the cutoff points determined previously [69]. Subject characteristics are summarized in Table S1.

### Plasma collection

At 9 AM, after an overnight fast from 10 PM the preceding evening, blood plasma was collected. Samples were immediately stored at −80 °C until further analysis.

### One-phase lipidomic extraction

In a 2 ml tube, the following amounts of internal standards dissolved in 1:1 (v/v) methanol: chloroform were added to each sample: BMP(14:0)_2_ (0.2 nmol), CL(14:0)_4_ (0.1 nmol), LPA(14:0) (0.1 nmol), LPC(14:0) (0.5 nmol), LPE(14:0) (0.1 nmol), LPG(14:0) (0.02 nmol), PA(14:0)_2_ (0.5 nmol), PC(14:0)_2_ (0.2 nmol), PE(14:0)_2_ (0.5 nmol), PG(14:0)_2_ (0.1 nmol), PS(14:0)_2_ (5 nmol), ceramide phosphocholine SM(d18:1/12:0) (2 nmol) (Avanti Polar Lipid). After adding the internal standard mix, a steel bead and 1.5 ml 1:1 (v/v) methanol: chloroform were added to each sample. Samples were homogenized using a TissueLyser II (Qiagen) for 5min at 30 Hz. Each sample was then centrifuged for 10 min at 20,000 *g*. The supernatant was transferred to a 4 ml glass vial.

### Lipidomics

The bottom layer, containing the apolar phase, was transferred to a clean 1.5 mL tube and evaporated under a stream of nitrogen at 60°C. The residue was dissolved in 100 μL of 1:1 (v/v) methanol:chloroform. Lipids were analyzed using a Thermo Scientific Ultimate 3000 binary HPLC coupled to a Q Exactive Plus Orbitrap mass spectrometer. For normal phase separation, 2 μL of each sample was injected onto a Phenomenex^®^ LUNA silica, 250 * 2 mm, 5µm 100Å. Column temperature was held at 25°C. Mobile phase consisted of (A) 85:15 (v/v) methanol:water containing 0.0125% formic acid and 3.35 mmol/L ammonia and (B) 97:3 (v/v) chloroform:methanol containing 0.0125% formic acid. Using a flow rate of 0.3 mL/min, the LC gradient consisted of: Dwell at 10% A 0-1 min, ramp to 20% A at 4 min, ramp to 85% A at 12 min, ramp to 100% A at 12.1 min, dwell at 100% A 12.1-14 min, ramp to 10% A at 14.1 min, dwell at 10% A for 14.1-15 min. For reversed phase separation, 5 μL of each sample was injected onto a Waters HSS T3 column (150 x 2.1 mm, 1.8 μm particle size). Column temperature was held at 60°C. Mobile phase consisted of (A) 4:6 (v/v) methanol:water and B 1:9 (v/v) methanol:isopropanol, both containing 0.1% formic acid and 10 mmol/L ammonia. Using a flow rate of 0.4 mL/min, the LC gradient consisted of: Dwell at 100% A at 0 min, ramp to 80% A at 1 min, ramp to 0% A at 16 min, dwell at 0% A for 16-20 min, ramp to 100% A at 20.1 min, dwell at 100% A for 20.1-21 min. MS data were acquired using negative and positive ionization using continuous scanning over the range of m/z 150 to m/z 2000. Data were analyzed using an in-house developed lipidomics pipeline written in the R programming language (http://ww.r-project.org). All reported lipids were normalized to corresponding internal standards according to lipid class, as well as to freeze-dried tissue weight. Lipid identification has been based on a combination of accurate mass, (relative) retention times, fragmentation spectra, analysis of samples with known metabolic defects, and the injection of relevant standards [70]. Lipidomics data from these experiments can be found in Table S3.

### Data analyses, statistics, and visualization

Data were processed and analyses were performed with R version 4.2.2 [71]. Data were processed in part with the R package dplyr version 1.1.4 [72], and tidyverse version 2.0.0 [73]. Significance was assessed using an empirical Bayes moderated t test on log2-transformed data within limma’s linear model framework, taking participant groups into account. Visualization of data was performed using ggplot2 version 3.4.4 [74], pheatmap version 1.0.12 [75], and VennDiagram version 1.7.3 [76].

## Author contributions

Conceptualization: W.L., G.E.J., R.H.H.; Human study design and collection of plasma: L.G., P.S., J.H., Formal analysis: W.L., B.V.S., G.E.J.; Investigation: W.L., A.W.G.; Data curation: B.V.S., G.E.J.; Writing - original draft: W.L., A.W.G. G.E.J., R.H.H.; Writing - review & editing: W.L., A.W.G. G.E.J., F.V., G.S.S., L.G., P.S., J.H., R.H.H.; Supervision: G.E.J., R.H.H.; Funding acquisition: G.E.J., R.H.H..

## Data availability

Lipidomics data are available as supplementary materials accompanying this manuscript as processed abundance values. All other data supporting the findings of this study are either also available as supplementary materials accompanying this manuscript or are available from the corresponding author upon reasonable request.

