## Supplemental table 1 and 2 for "Plasma triacylglycerol length and saturation level mark healthy aging groups in humans"

Table 1. Cohort characteristics.

|  |  | Older adults | | |
| --- | --- | --- | --- | --- |
|  | Young | Trained | Normal | Impaired |
| Participants | 14 | 19 | 16 | 3 |
| Age range (yr) | 20-30 | 65-80 | 65-80 | 65-80 |
| Male (%) | 50% | 58% | 50% | 67% |
| BMI (kg m-2) | 22.63 | 23.64 | 25.58 | 26.30 |
| Steps per day | 10359 | 13815 | 9986 | 7290 |
| Exercise (/week) | <1h | >3h | <1h | <1h |
| Time high activity (%) | 2.81 | 5.34 | 2.17 | 1.22 |

Table 2. List of lipids detected in participants.

| **Category** | **Main class** | **Class key** | **Common name** | **Nb detected** |
| --- | --- | --- | --- | --- |
| **Glycerolipids** | Diacylglycerols | DG | Diacylglycerol | 169 |
|  | Triacylglycerols | TG | Triradylglycerol | 348 |
|  |  | TG[O] | Alkyldiacylglycerol | 90 |
| **Glycerophospholipids** | Glycerophosphocholins | LPC | Lysophosphatidylcholine | 63 |
|  |  | LPC[O] | Alkyllysophosphatidylcholine | 41 |
|  |  | PC | Phosphatidylcholine | 108 |
|  |  | PC[O] | Alkylphosphatidylcholine | 98 |
|  | Glycerophosphoethanolamines | LPE | Lysophosphatidylethanolamine | 20 |
|  |  | LPE[O] | Alkyllysophosphatidylethanolamine | 19 |
|  |  | PE | Phosphatidylethanolamine | 39 |
|  |  | PE[O] | Alkylphosphatidylethanolamine | 41 |
|  | Glycerophosphoinositols | PI | Phosphatidylinositols | 19 |
|  | Glycerophosphates | LPA | Lysophosphatic acid | 11 |
|  |  | PA | Phosphatidic acid | 14 |
| **Sphingolipids** | Phosphosphingolipids | SM[d] | Sphingomyelin | 82 |
|  |  | SM[t] | Hydroxysphingomyelin | 36 |
|  | Acidic glycosphingolipids | SM4[d] | Sulfatide | 12 |
|  |  | SM4[t] | Hydroxysulfatide | 15 |
|  | Ceramides | C1P[d] | Ceramide-1-phosphate | 8 |
|  |  | Cer[d] | Ceramide | 69 |
|  | Neutral glycosphingolipids | Hex2Cer[d] | Dihexosylceramide | 17 |
|  |  | HexCer[d] | Hexocylceramide | 28 |
|  | Sphingoid bases | S1P | Sphingosine | 4 |
| **Sterol lipids** | Sterols | CE | Cholesteryl ester | 95 |
